# Antibodies targeting a conserved cryptic epitope at the influenza hemagglutinin head-stem interface via distinct binding modes

**DOI:** 10.64898/2026.09.24.754278

**Authors:** Guthrie L. Stroh, Huibin Lv, Chunke Chen, Yuanxin Sun, Tossapol Pholcharee, Catherine Zhu, Shu Ting Teo, Douglas A. Mitchell, Chris K.P. Mok, Nicholas C. Wu

**Affiliations:** Department of Chemistry, University of Illinois Urbana-Champaign, Urbana, IL 61801, USA; Department of Biochemistry, University of Illinois Urbana-Champaign, Urbana, IL 61801, USA; Carl R. Woese Institute for Genomic Biology, University of Illinois Urbana-Champaign, Urbana, IL 61801, USA; The Jockey Club School of Public Health and Primary Care, The Chinese University of Hong Kong, Hong Kong SAR, China; Li Ka Shing Institute of Health Sciences, Faculty of Medicine, The Chinese University of Hong Kong, Hong Kong SAR, China; Department of Microbiology, Faculty of Science, Mahidol University, Bangkok 10400, Thailand; Department of Biochemistry, Vanderbilt University School of Medicine—Basic Sciences, Nashville, TN, USA; Department of Chemistry, Vanderbilt University, Nashville, TN, USA; S. H. Ho Research Centre for Infectious Diseases, The Chinese University of Hong Kong, Hong Kong, China; School of Biomedical Sciences, The Chinese University of Hong Kong, Hong Kong SAR, China; Center for Biophysics and Quantitative Biology, University of Illinois Urbana-Champaign, Urbana, IL 61801, USA; Carle Illinois College of Medicine, University of Illinois Urbana-Champaign, Urbana, IL 61801, USA

## Abstract

Influenza A virus continues to pose pandemic threats, as demonstrated by the recent global spread of highly pathogenic H5N1 strains. Consequently, the development and characterization of vaccines targeting zoonotic influenza subtypes have become a major focus of the field. In this study, we characterize two broadly reactive antibodies previously isolated from H5N1 vaccinees. Although they have distinct immunoglobulin germline gene usage and binding modes, cryo-EM analysis shows that both antibodies share a highly conserved cryptic epitope at the interface of the hemagglutinin (HA) head and stem domains. The antigenicity of this epitope is influenced by natural amino acid variation at residue 293 of HA subunit 1, and its access is facilitated by acid-induced conformational change of HA. Nevertheless, both antibodies are non-neutralizing and possess only marginal protective efficacy against seasonal and H5N1 viruses *in vivo*, thus representing suboptimal vaccine responses. Overall, our findings expand the interactions of cross-reactive antibodies with an under-characterized HA epitope and provide insights for next-generation influenza vaccine design.

## INTRODUCTION

Global circulation of influenza virus among the human population has played out on the cyclical stage of host immune response and antigenic evolution. The majority of molecular divergence among influenza viruses occurs in the two predominant surface glycoprotein antigens, hemagglutinin (HA), responsible for viral attachment, and neuraminidase (NA), facilitating viral release.^1^ Alongside influenza B virus, eight HA subtypes of influenza A virus have to date demonstrated an ability to infect humans: H1, H2, H3, H5, H6, H7, H9, and H10; an additional ten subtypes are have thus far been found exclusively in animal reservoirs (H4, H8, and H11-18).^2^ Seasonal influenza vaccines focus on stimulating humoral response against H1N1 and H3N2 subtypes of influenza A virus and the phylogenetically distant influenza B virus, since they currently circulate in humans and cause annual epidemics.^3,4^

Influenza vaccination or infection typically elicits antibodies that recognize the highly variable head domain of HA, resulting in a lack of a strong heterosubtypic antibody response.^5,6^ Therefore, much of the work in the influenza field over the past decade has centered around achieving a vaccination capable of eliciting broadly protective antibodies capable of binding diverse HAs.^7,8^ Defense against the highly pathogenic H5N1 strain of influenza A virus represents a critical portion of this effort, as recent outbreaks of mammalian-adapted H5N1 virus among livestock have showcased the threat of zoonotic spillover, potentially triggering a human pandemic.^9,10^ Many antibodies with the ability to recognize a broad range of HA subtypes, including H5, have been reported.^11^ These antibodies target highly conserved epitopes on HA, such as the receptor binding site (RBS) and trimer interface in the head domain, as well as multiple epitopes in the stem domain.^12–18^ Given that many of these broad antibodies are also capable of protecting against lethal influenza virus infection, an understanding of their interactions with HA is critical for developing broadly protective influenza vaccines.

Immunogen design is a key facet of the development of broadly protective influenza vaccines.^19^ A growing body of evidence suggests that steric accessibility of different HA epitopes is a central driver of their immunodominance hierarchies.^20,21^ Efforts in eliciting broadly reactive antibodies have been successful by presenting HA stem-only protein constructs^22^, using antigens which invert the HA trimer to facilitate access to its stem^23^, utilizing mosaic HA nanoparticles^24^, or by leveraging computationally optimized consensus sequence antigens that provide reactivity across HA strains.^25^ Recently, an increasing number of vaccination studies have been conducted with a primary focus on H5N1 strains^26–30^ and a thorough examination of their immunological outcomes will be critical in driving the rational development of future broadly protective vaccines.

Here, we characterized two broadly reactive human antibodies previously isolated from experimental H5N1 vaccinees,^31^ both of which bound to an under-characterized yet highly conserved HA epitope at the interface between its head and stem domains. While this cryptic epitope conferred broad reactivity, both antibodies were shown to be non-neutralizing and only weakly protective *in vivo*. Structural analysis revealed that these two antibodies had distinct binding modes. Subsequent mutagenesis experiments further highlighted a key residue of this epitope governing its antigenicity. Taken together, the work presented here will be a valuable resource in refining the development of broadly protective influenza immunogens.

## RESULTS

### Discovery of two broadly reactive, non-stem-binding antibodies

We have recently reported the binding profile for over 300 publicly available and natively paired influenza HA monoclonal antibodies (mAbs)^32^. Many antibodies from this study showed heterosubtypic binding breadth, which is well-documented for known conserved HA epitopes such as HA stem,^33^ HA RBS,^34^ and HA trimer interface.^35^ To uncover novel epitopes that can be recognized by cross-reactive HA antibodies, we narrowed our focus by removing mAbs with common germline gene usage or well-characterized sequence features in their complementarity-determining regions (CDRs). Additionally, mAbs that did not bind to the H1 stem^36^ and H3 stem^37^ were prioritized. Under these selection criteria, two mAbs, 01.z.01 and 56.e.01, which were isolated from two different human individuals in a clinical trial of an H5N1 vaccine (VRC 310), were prioritized for further investigation.^31,38^

Both 01.z.01 and 56.e.01 bound to many diverse influenza A HA subtypes from both group 1 and group 2 via the enzyme-linked immunosorbent assay (ELISA) (**Figure 1A-B)**. However, neither 01.z.01 nor 56.e.01 had any detectable neutralization activity to representative H1N1 and H3N2 viruses as measured by the hemagglutination inhibition assay (HAI) (**Figure 1C**) and microneutralization assay (MN) (**Figure 1D**). This non-neutralizing property has been reported for other mAbs that recognize epitopes on HA that appear buried in the prefusion trimer.^17,35,39^ To identify the epitopes targeted by 01.z.01 and 56.e.01, we designed a competition assay using the two cross-reactive antibodies 5J8^40^ and FluA-20,^17^ which are known to bind the RBS and trimer interface, respectively (**Figure 1E)**. We also included S8V1-157, which was reported during the course of the current study and binds to the interface between the HA head and stem domains.^41^ In this all-by-all biolayer interferometry (BLI) competition screen, 01.z.01, 56.e.01, and S8V1-157 Fabs all competed with each other for binding (competition index > 0.5), but neither 01.z.01 nor 56.e.01 competed with either FluA-20 or 5J8 (**Figure 1F**). These data indicated that both 01.z.01 and 56.e.01 bind to or near the recently identified head-stem interface epitope.^41^ Notably, 56.e.01 possessed similar features to other S8V1-157-competing mAbs, including germline usage of *IGKV2-28*01* and *IGHJ4*02*, alongside a “MQALQ” motif in its LCDR3 which mirrored contacts of S8V1-157 with HA^41^ (**Figure 1G**). By contrast, 01.z.01 lacked all these features.

**Figure 1.**
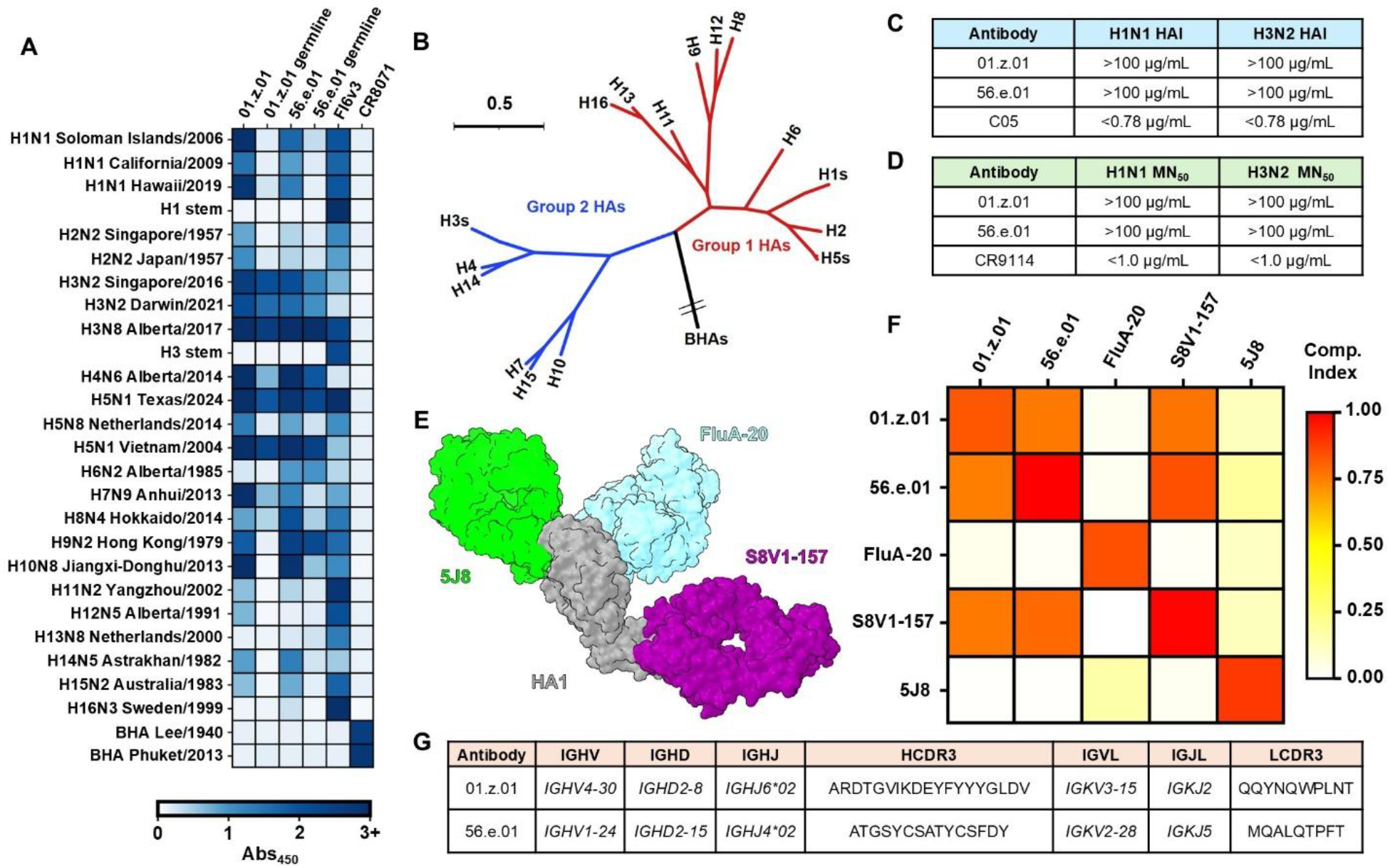
01.z.01 and 56.e.01 are broadly binding and non-neutralizing antibodies. **(A)** The HA binding profile of 01.z.01 and 56.e.01, alongside their germline revertants, was determined by ELISA. Plotted heatmap values show the absorbance at 450 nm averaged across two replicates. Full HA strain names can be found in Figure 3D. **(B)** Phylogenetic tree depicting the primary amino acid sequence divergence between HA subtypes used in this study. **(C)** *In vitro* activity of 01.z.01 and 56.e.01 IgGs as measured by the hemagglutination activity inhibition (HAI) assay against H1N1 A/California/04/2009 or H3N2 A/Philippines/2/1982 virus. Antibody C05^84^ is used as a positive control. **(D)** *In vitro* activity of 01.z.01 and 56.e.01 IgGs as measured by the microneutralization (MN) assay against H1N1 A/California/04/2009 or H3N2 A/Philippines/2/1982 virus. Antibody CR9114^15^ is used as a positive control. **(E)** Structural representation of the three Fabs used in the binding competition assay. 5J8 (green, RBS, PDB 4M5Y),^85^ FluA-20 (light blue, trimer interface, PDB 6OC3),^17^ and S8V1-157 (purple, head-stem interface, PDB 8US0)^41^ cover all known highly conserved epitopes on the HA head (grey). **(F)** Competition indices from the biolayer interferometry (BLI) binding assay. Heatmap values represent degree of binding competition on A/California/04/2009 (H1N1) head domain, from mutually inclusive (white) to mutually exclusive (red). **(G)** Germline gene usage and CDR3 sequences of 01.z.01 and 56.e.01.

### 01.z.01 targets the head-stem interface epitope with a novel binding mode

To elucidate the molecular recognition of HA by 01.z.01, we determined the structure of HA head domain of H3N2 A/Darwin/9/2021 (H3/Darwin21) in complex with 01.z.01 Fab, FluA-20 Fab,^17^ and ADI-85647 Fab^42^ using cryogenic electron microscopy (cryo-EM). The inclusion of FluA-20 Fab and ADI-85647 Fab, whose epitopes do not overlap with the hypothesized epitope of 01.z.01, served to increase the molecular weight of the complex to facilitate cryo-EM analysis. This quadripartite complex remained stable during size-exclusion chromatography and yielded a cryo-EM map with a global resolution of 3.38 Å and a local resolution of 3.0 Å at the 01.z.01-HA interface (**Table S1, Figure S1A-D, left**).

The epitope targeted by 01.z.01 was revealed to be highly occluded, lying nearly completely buried in the prefusion conformation of HA trimer (**Figure 2A, left**). The epitope was further shielded by the intra-protomer association of the C-terminus of HA1 and the N-terminus of HA2 (**Figure 2A, middle**). Residing on the bottom of the HA head domain, the 01.z.01 epitope was only fully exposed upon dissociation of the stem domain, a process that occurs under acidic conditions to promote virus-host membrane fusion^43^ (**Figure 2A, right**). While the epitope of 01.z.01 partially overlapped with that of S8V1-157,^41^ we found that there were several key differences that distinguished 01.z.01 and explained its absence of shared sequence features. Firstly, 01.z.01 has an entirely different angle of approach to the head-stem interface, with approximately 90° of axial HA rotation separating its approach from that of S8V1-157 (**Figure 2B-C**). 01.z.01 also had an interaction interface with a total buried surface area (BSA) of 1052.2 Å^2^, which was 9.2% larger than that of S8V1-157 (**Figure S2**). This disparity is elucidated by the presence of 01.z.01 contacts spanning HA residues N- and C-terminal to the border of the S8V1-157 epitope (**Figure 3D**). The 01.z.01 epitope was mainly comprised of HA1 residues that are highly conserved across H1-H16 subtypes, which rationalized the expansive breadth of its HA recognition (**Figure 2E**). This level of sequence conservation can be explained by the functional importance of stable intramonomer sidechain packing during protein folding, along with the relatively steep mutational fitness burden of residues under multiple epistatic interaction constraints.^44–46^ Additionally, this region of HA has likely experienced little immune selection pressure, as it is buried in the prefusion conformation, unlike frequently targeted and thus varied regions on the solvent-exposed surface of the HA head domain.^47^

**Figure 2.**
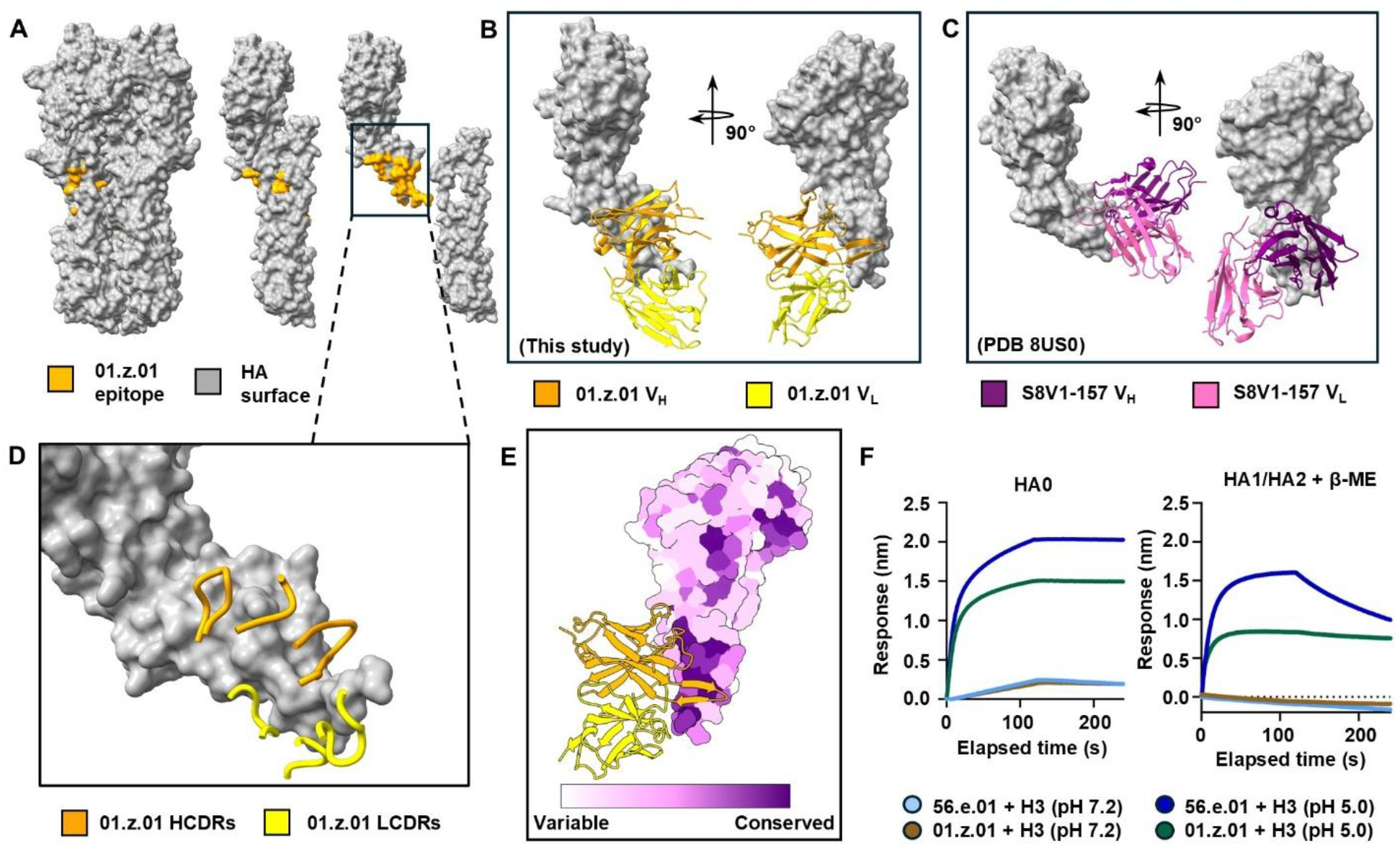
01.z.01 binds to a conformationally sensitive and highly conserved cryptic epitope at the HA head-stem interface with a ‘side-on’ approach angle. **(A)** The 01.z.01 epitope (orange) is partially shielded in trimeric and monomeric HA (grey, left and middle). The epitope is fully revealed upon the dissociation of the HA head and stem (right). **(B)** 01.z.01 scFv (V_H_: orange, V_L_: yellow) binds to A/Darwin/9/2021 (H3N2) HA (grey) with 90 degrees of axial rotation relative to the interface of the head and stem domains. **(C)** S8V1-157 scFv (PDB 8US0, V_H_: purple, V_L_: pink)^41^ binds to A/American black duck/New Brunswick/00464/2010 (H4N6) HA (grey) in a ‘head-on’ angle of approach relative to the interface of the head and stem domains. **(D)** Close up extension of the 01.z.01 epitope from panel **A**, depicting the orientation of the 01.z.01 HCDRs (orange) and LCDRs (yellow) on the HA surface (grey). **(E)** 01.z.01 scFv (orange, yellow) engages highly conserved residues at the head-stem interface. The HA surface is colored according to residue conservation scores produced from the sequence alignment depicted in Figure 3D. **(F)** Biolayer interferometry binding response of immobilized 01.z.01 and 56.e.01 Fabs to 300nM of full-length HA ectodomain from A/Darwin/9/2021 (H3N2) after extended incubation under neutral or acidic pH. Shown are trimeric uncleaved HA0 (left) and HA1/HA2 produced after trypsinolysis and reduction with β-mercaptoethanol (β-ME) (right).

**Figure 3.**
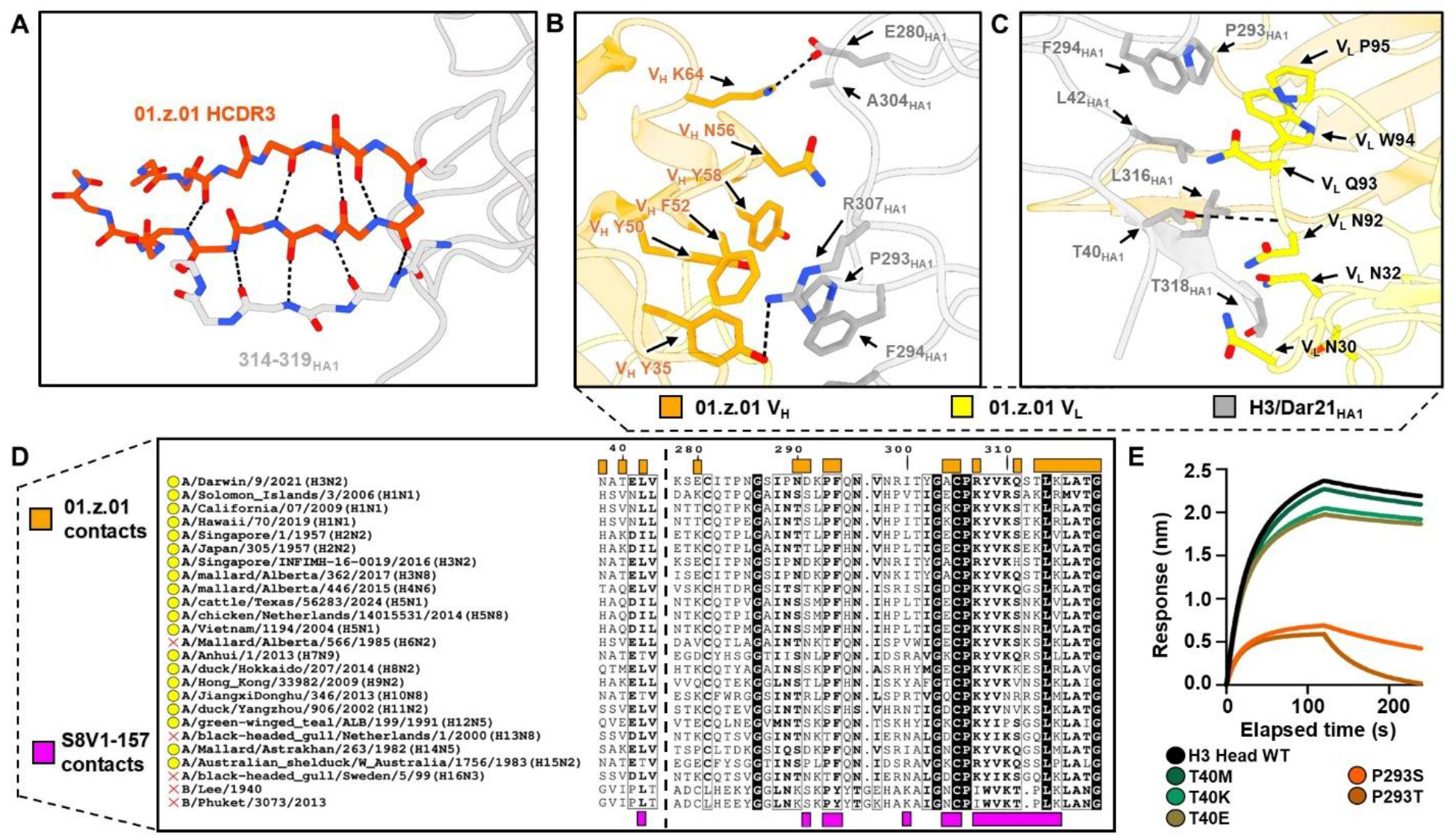
Molecular analysis and site-directed mutagenesis of the 01.z.01 HA epitope reveal a key breadth-determining position. **(A)** The 01.z.01 HCDR3 (orange) makes multiple continuous backbone-backbone contacts (black dashed lines) to form an extended β-strand with HA1 residues 314-319 (light grey, H3 numbering). **(B**,**C)** Molecular recognition of HA (grey) by 01.z.01 V_H_ (orange, left) and 01.z.01 V_L_ (yellow, right). Hydrogen bonding interactions (black dashed lines) are shown. **(D)** Sequence alignment of the HA strains used in this study. Sequences are abbreviated to depict HA contacts with 01.z.01 (orange boxes) and S8V1-157 (purple boxes). Strains bound by 01.z.01 in ELISA are labeled with a yellow circle, whereas those 01.z.01 failed to bind have a red cross. **(E)** Biolayer interferometry binding curves for 500 nM 01.z.01 Fab against a panel of immobilized A/Darwin/9/2021 (H3N2) HA1 single-site variants.

To gain insight into the accessibility of the head-stem interface epitope, we tested the binding of 01.z.01 or 56.e.01 to the uncleaved H3/Darwin21 HA0 at neutral pH or after a 15-min exposure to pH 5.0, which mimicks the acidic conditions during viral fusion.^43^ While the two antibodies showed weak binding to the uncleaved HA0 trimer at neutral pH, there was a marked increase of binding signal after the acidic incubation (**Figure 2F**). When the same H3/Darwin21 HA was cleaved into HA1/HA2 and HA1 was liberated via reduction of the interchain disulfide, the binding response of both 01.z.01 and 56.e.01 was only detected upon acidic incubation (**Figure 2F**). After quantifying the binding of both Fabs to head-only H1 and H3 constructs, we found 01.z.01 had a K_d_ of 23 nM and 24 nM to H1 and H3 heads, respectively, which was comparable to other protomer interface Fabs.^17,41,48^ However, 56.e.01 possessed notably weaker affinity, with a K_d_ of 213 nM to H1 and 55 nM to H3 (**Figure S3**). Taken together, these results indicate that the association of 01.z.01 and 56.e.01 during the viral life cycle is likely dictated by the relationship between differential epitope accessibility and prefusion conformational stability of HA.^49,50^

### Analysis of important residues for 01.z.01 HA binding affinity

The CDRs of 01.z.01 wrap around a protruding end of the HA head domain comprised of the N and C-termini of HA1 (**Figure 2D**). The 19-residue HCDR3 of 01.z.01, which accounted for 36% of the total paratope BSA, adopted an intramolecular β-hairpin motif that interacted with residues 314-319 of HA1 (H3 numbering) to form an extended parallel β-sheet (**Figure 3A and Figure S2**). Because the formation of this β-sheet relies entirely on backbone-backbone interactions, differences at these residues among diverse HA strains retain binding to 01.z.01 (**Figure 3D**). Structural analysis of 01.z.01 beyond the HCDR3 revealed the majority of the paratope was comprised of relatively apolar residues packed against a similarly hydrophobic HA interaction surface (**Figure 3B-C**). Only two specific sidechain-sidechain interactions were observed, with V_H_ K64 (Kabat numbering) forming a salt bridge with E280_HA1_ and V_H_ Y35 H-bonding with the guanidinium group of R307_HA1_ (**Figure 3B-C**). Notably, at residue 307_HA1_, group 1 and 2 HAs typically possess Lys or Arg, whereas influenza B HAs possess an Ile (**Figure 3D**), potentially explaining the absence of 01.z.01 binding to influenza B HAs (**Figure 1A**). P293_HA1_, which interacted with both V_H_ and V_L_, stood out as a less conserved epitope residue, with H11 having a Ser at this residue, while H6, H13, and H16 carry a Thr (**Figure 3D**). Variant P293S_HA1_ in H3/Darwin21 drastically reduced the *k*_*on*_ of 01.z.01, and P293T_HA1_ further increased the *k*_*off*_ (**Figure 3E**). This mutational data aligned well with the ELISA binding, as H6, H13, and H16 were the only HA subtypes to show no significant binding (**Figure 1A**). Importantly, S8V1-157 showed a similar binding reduction to the P293S_HA1_ variant and binding to P293T_HA1_ was entirely abrogated, which corroborated the reported ELISA binding breadth of S8V1-157 (**Figure S4**).^41^ Therefore, P293_HA1_ is a key breadth determinant of antibodies targeting the head-stem interface epitope with distinct binding modes.

The HA residue with the lowest conservation on the 01.z.01 epitope was T40_HA1_, the hydroxyl of which was predicted to form a hydrogen bond to the backbone of V_L_ N92 (**Figure 3C**). While HA residue 40 has multiple physicochemically diverse amino acid variants across subtypes, mutating T40_HA1_ to other naturally existing amino acid variants Glu, Met, and Lys had no substantial impact on the binding of 01.z.01 to HA (**Figure 3E**). S8V1-157 also retained strong binding to these variants, likely because this residue lies outside of its binding footprint (**Figure S4**). While this latter discovery supported the robustness of molecular recognition by 01.z.01, it also indicated that disparities among the binding activity of 01.z.01 to P293_HA1_-possessing subtypes are likely due to higher-order interaction networks that are not easily discerned from sequence or structural alignments. For example, while the salt bridge between V_H_ K64 and E280_HA1_ might appear important for binding, E280 is poorly conserved among group 1 HAs such as H5, which demonstrated binding to 01.z.01 despite having a non-charge-complementary Lys at this position (**Figure 1A, Figure 3D**).

We then probed the importance of somatic hypermutation (SHM) on the binding breadth of 01.z.01. As shown by ELISA, the breadth of 01.z.01 was narrower upon germline reversion (**Figure 1A, Figure S5A**). The binding of 01.z.01 germline revertant to H5 aligned with isolation from H5N1 vaccinees, yet it completely lost binding to the antigenically closer H1 and H2 HAs, while retaining strong binding to the much more distantly related H3 HAs (**Figure 1A-B**). SHMs V_H_ S32D on the HCDR1 and V_H_ Y53H on the HCDR2 form an intramolecular ionic interaction (**Figure S5A-B**). This salt bridge may pre-organize and stabilize the aromatic pocket of V_H_ Y35 and V_H_ F52, both of which were directly implicated in binding the charge-conserved R307_HA1_ (**Figure S5B**). Many of the remaining germline SHMs were either chemically similar residues or existed at locations distal from the paratope (**Figure S5A & C**). While we could not readily interpret such contributions, multiple studies have demonstrated the importance of such non-paratope SHMs in breadth expansion.^51–55^ In total, the 11.1% and 5.5% V_H_ and V_L_ SHM burden, respectively, indicated 01.z.01 underwent extensive affinity maturation to arrive at its broad HA binding profile.

### 56.e.01 shares a binding footprint with previously characterized mAbs

To characterize the epitope of 56.e.01, we determined a low-resolution structure of H3/Darwin21 HA head domain in complex with 56.e.01 Fab, FluA-20 Fab, and ADI-85647 Fab. (**Table S1, Figure S1A-D, right**). We then fitted an AlphaFold3-predicted model^56^ of the variable domain of 56.e.01 in complex with H3/Darwin21 head domain into the resulting 8.8 Å map. 56.e.01 appeared to engage the head-stem interface epitope with a similar binding footprint and approach angle as S8V1-157 (**Figure 4A-B**). The predicted orientation of the LCDRs in 56.e.01 and those of S8V1-157 are nearly identical, consistent with their shared light chain sequence features (**Figure 4C**). One notable difference came from the HCDR3 of 56.e.01, predicted to contain a disulfide bond (*C*SATY*C*), which, in our model, occupied a similar region on HA to the HCDR2 of S8V1-157. Additionally, the HCDR2 model of 56.e.01 was shifted out and away from the surface of HA, seemingly making minimal direct contacts (**Figure 4C**). This latter feature may explain the binding affinity disparity between 56.e.01 and other head-stem interface mAbs.

**Figure 4.**
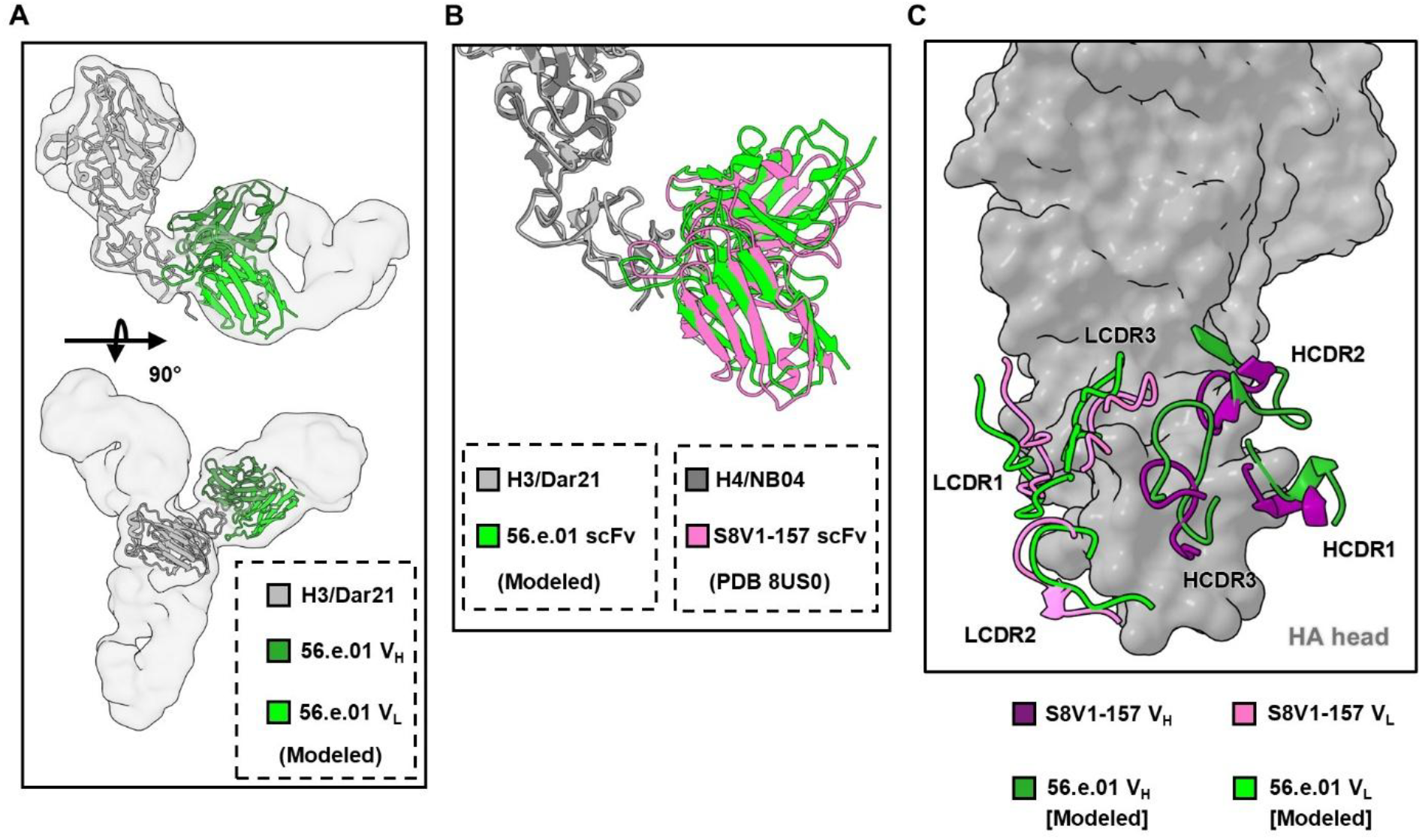
Analysis of the low-resolution 56.e.01 + H3 head complex and comparison to previously reported head-stem interface antibody S8V1-157. **(A)** The AlphaFold3^56^ model of 56.e.01 scFv (V_H_: dark green, V_L_: light green) in complex with A/Darwin/9/2021(H3N2) HA1 (light gray) fitted into its respective cryo-EM map. **(B)** Structural alignment of the 56.e.01 scFv (light green) + H3 head (light gray) model with the S8V1-157 scFv (pink) + A/American black duck/New Brunswick/00464/2010 (H4N6) (dark gray) complex, adapted from PDB 8US0.^41^ Alignment was calculated using HA backbone atoms. **(C)** Comparison of the CDR positioning on the HA surface (gray) between the 56.e.01 model and S8V1-157.

When we performed the germline reversion experiments with 56.e.01, a similar pattern of breadth contraction to 01.z.01 was observed (**Figure 1A**). 56.e.01 germline revertant also retained H3 238 and H5 binding, while H1 binding was notably lost. The one significant disparity between the germline revertants was that only 56.e.01 germline revertant retained strong H9 binding (**Figure 1A**). These results demonstrated that antibodies targeting the head-stem interface epitope with distinct binding modes could originate from germlines with similar HA binding specificities.

### 01.z.01 and 56.e.01 have weak *in vivo* protective activity

To assess the protective efficacy of 01.z.01 and 56.e.01, we expressed both antibodies under the Fc backbone of mouse IgG1 (mIgG1) or mouse IgG2c (mIgG2c). While mIgG1 possesses tissue inflammation-inhibiting activity through potent FcγRIIB (CD32B) binding, it drives little complement activation.^57,58^ By contrast, mIgG2c binds strongly to multiple activating FcγRs, potently eliciting effector functions including antibody-dependent cellular cytotoxicity, antibody-dependent cellular phagocytosis, and the C1q complement cascade.^59–61^ Therefore, studying both mIgG subclasses in tandem would provide insight into the reliance of 01.z.01 and 56.e.01 on Fc-dependent protective mechanisms. All mice prophylactically treated with 10 mg/kg of 01.z.01 mIgG1, 56.e.01 mIgG1, or 56.e.01 mIgG2c succumbed to a lethal intranasal challenge of H1N1 A/California/04/2009 (H1/Cal09), whereas 20% of mice treated with the same dose of 01.z.01 mIgG2c survived (**Figure 5A, 5C, 5E**, & **5G**). This Fc-dependent effect was more apparent when the mice received a lethal challenge of H3N2 A/Philippines/2/1982 (X-79), in which case 01.z.01 conferred 20% and 100% survival rates when formatted as mIgG1 and mIgG2c, respectively (**Figure 5B & 5D**). In contrast, the Fc-dependent effect was less pronounced for 56.e.01, which conferred 20% and 40% survival rates against X-79 as mIgG1 and mIgG2c, respectively (**Figure 5F & 5H**). Consistently, the lung viral titers of X-79-challenged mice at day 3 post-infection showed a nearly two-log reduction when 01.z.01 was administered as mIgG2c compared to mIgG1 (p = 0.02), but such difference was insignificant for 56.e.01 (**Figure 5K**). The disparity between 01.z.01 and 56.e.01 in Fc-dependent protection may be due, at least in part, to the difference in their approach angles to the head-stem interface epitope (**Figure 2B, Figure 4A**), which in turn may influence their access to Fc receptors.^62,63^

**Figure 5.**
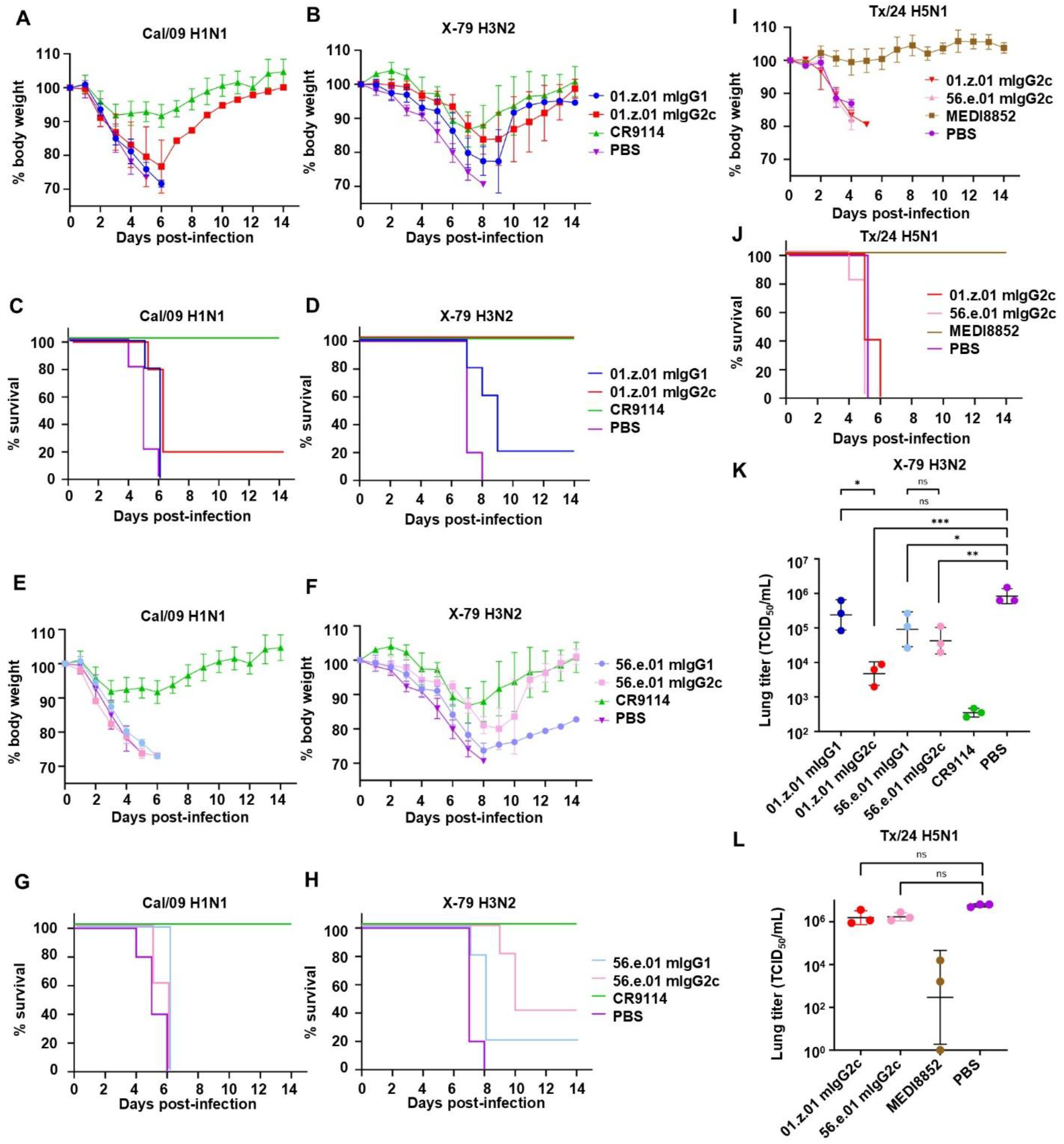
01.z.01 and 56.e.01 have poor *in vivo* protective efficacy. Female BALB/c mice at six weeks old were intraperitoneally injected with 10 mg/kg of the indicated antibody (or 5 mg/kg for CR9114) 4 h prior to intranasal challenge with 5× LD50 of H3N2 A/Philippines/1982 (X-79), H1N1 A/California/04/2009 (Cal/09), or H5N1 A/Texas/37/2024 (Tx/24). In the case of H5N1 challenge, mice were instead injected with 25 mg/kg of the indicated antibodies. **(A-B, E-F & I)** Mean percentage of initial body weight post-infection is depicted over the 14-day course of the experiment (n = 5 for each experimental and control group). The humane endpoint of a 25% decrease from starting body weight was instituted. Data are presented as mean ± standard deviation. **(C-D, G-H & J)** Kaplan-Meier survival curves for each treatment and control group (n = 5 each) in each viral challenge condition are presented. Curves are slightly offset for clarity. **(K-L)** Three-day lung viral titers in median tissue culture infectious dose (TCID_50_) for each treatment and control group (n = 3 each) for X-79 **(K)** or Tx/24 **(L)** are shown. Each graphed point represents the average across three technical replicates for one mouse. Data are presented as geometric mean ± geometric standard deviation. * = p < 0.05; ** = p < 0.01; *** = p < 0.001; ns = not significant (p > 0.05). P-values were determined using a homoscedastic one-tailed Student’s t-test of the log_2_-transformed TCID_50_ values.

Since 01.z.01 and 56.e.01 were isolated from H5N1 vaccinees, we next evaluated their protective efficacy against the H5N1 clade 2.3.4.4b strain A/Texas/37/2024 (Tx/24) which was isolated from a human patient^64^. Importantly, both mAbs showed strong binding activity to Tx/24 H5 (**Figure 1A**). Given the highly pathogenic nature of H5N1, the extreme virulence of which allows rapid disease progression to multiple vital organs,^64^ we increased the prophylactic mAb dose to 25 mg/kg to amplify the likelihood of detecting protective activity. Despite this high dose, neither 01.z.01 mIgG2c nor 56.e.01 mIgG2c provided protection against lethal Tx/24 H5N1 challenge as measured by the weight loss, survival rate, and lung viral titers at 3 days post-infection (**Figure 5I-J & 5L**). In contrast, all mice treated with 25 mg/kg of central stem antibody MEDI8852^52^ survived with almost no weight loss (**Figure 5I-J**) and exhibited at least a two-log reduction of lung viral titer at 3 days post-infection compared to the untreated mice (**Figure 5L**). Together, these data indicated that 01.z.01 and 56.e.01 had suboptimal protective efficacy *in vivo*.

## DISCUSSION

The pandemic threat of influenza A virus has motivated the development of novel immunogens that are protective against both seasonal infections and pathogenic zoonotic subtypes. Such immunogen development requires a rigorous structural and functional understanding of the antibodies they elicit, especially in human trials. In this study, we characterized two HA antibodies, 01.z.01 and 56.e.01, both of which were isolated from a clinical trial of an experimental H5N1 vaccine.^31^ Both 56.e.01 and 01.z.01 had a broad HA recognition profile and engaged a recently discovered and highly conserved cryptic epitope at the interface of the head and stem domains. Although 56.e.01 had a binding mode similar to the only previously known antibody targeting this head-stem interface epitope,^41^ the binding mode of 01.z.01 was unique. Both 01.z.01 and 56.e.01 were non-neutralizing and conferred only weak protective activity *in vivo*. Overall, the results of this study provide insights into the optimization of HA-based immunogen design.

56.e.01 possessed a shared set of germline genes and LCDR motifs to a class of antibodies previously reported to target the head-interface epitope.^41^ Combined with the low-resolution map of 56.e.01 bound to HA presented here, it is evident that 56.e.01 belongs to this previous antibody class, albeit also expanding the known diversity of HCDR3 sequences capable of engaging the head-stem interface epitope in a ‘head-on’ angle of approach. In contrast, 01.z.01 had a ‘side-on’ approach angle to this epitope, with HA contacts outside of the previously defined interface. Despite this difference in angle of approach, both 01.z.01 and 56.e.01 retained a similar HA binding breadth upon germline reversion, demonstrating that disparate antibody lineages with distinct structural features remain subject to nearly identical constraints on head-stem interface antigenic recognition. In this case, the lineages of 01.z.01 and 56.e.01 both appear to arise from H3 or H5 immunization, with subsequent evolutionary trajectories converging on the broad influenza A recognition profile demonstrated here.

A notable structural feature of the 01.z.01-HA head complex was the orientation of the 19-residue HCDR3; all but two side chains faced away from the HA epitope. Instead, the HCDR3, which adopted a β-hairpin conformation, made multiple continuous backbone-backbone hydrogen bonds with HA, imparting an adopted β-strand structure on an otherwise tightly packed loop of HA1 (**Figure 3A**). This protein-protein interaction paradigm represents an addition to the current structural repertoire of known HCDR3-HA interactions and may account for some of the binding breadth observed in 01.z.01. It is likely length, rather than residue identity, that constrains HCDR3 compatibility with side-on head-stem interface recognition, suggesting that 01.z.01-like antibodies could be rather common among human lineages. Notably, one of the 11 mAbs previously reported to compete with S8V1-157 not only used similar germline genes (*IGHV4-31*04, IGVK3-15*01*, and *IGHJ6*02*), but also possessed a 19-residue HCDR3, making this mAb (S12V4_P1-F1) a strong contender for sharing this newly disclosed head-stem interface epitope approach angle and a good candidate for future study.^41^

One key finding of this study was the relatively poor protective efficacy of 01.z.01 and 56.e.01. In pursuit of a more universal vaccine design, protective efficacy is crucial. Given the recent clinical successes of broadly neutralizing antibodies against the HA stem,^65^ it is possible that antibodies recognizing and inhibiting a functional epitope on HA may fare better in future clinical development. This latter point is illustrated by our finding that against three viruses tested, including the highly pathogenic H5N1, only one head-stem interface mAb group (01.z.01-mIgG2c against H3N2 X-79) saw complete protection, and these antibody treatments were outperformed by central HA stem antibodies CR9114^15^ or MEDI8852.^52^ However, in the case of 56.e.01, it is likely that low binding affinity governed the weak protective activity, as several other mAbs binding the same epitope provided robust H3N2 protective activity.^41^ Given the high affinity of 01.z.01 for H1, H3, and H5, the weak protective activity may instead arise from the unique approach angle and thus Fc domain orientation compared to other known head-stem interface mAbs with stronger protective activity, at least against H3N2, such as S8V1-157 (**Figure 2B and Figure 2C**). Future studies interrogating the impact of head-stem interface mAb Fc angle may provide valuable mechanistic insight into the process of FcR- and C1q-mAb recognition for this epitope, as Fc function has been shown to influence the efficacy of mAbs to both binding modes.^41^ In fact, mAbs binding other occluded or conformationally-restricted epitopes often possess a similarly high reliance on Fc effector function for protective efficacy.^35,66–68^ Thus, the precise mechanism by which such mAbs, including 01.z.01 and 56.e.01, inhibit viral replication and spread in the host ought to be investigated in future studies. Given that even an exceedingly high dose of either 01.z.01 or 56.e.01 resulted in no detectable protective phenotype against H5N1, such future investigations would benefit from a critical evaluation of the role protomer interface-binding mAbs have in defending against the highly virulent, mammalian-adapted H5N1 strains posing a current zoonotic threat.

01.z.01 and 56.e.01 were isolated from subjects 1 and 56, respectively, out of six subjects in a previous study.^31^ The presence of these non-neutralizing antibody responses in subjects 1 and 56 may help explain why their fold increases of sera neutralization titers after vaccination were among the lowest in this cohort.^31^ As 01.z.01 and 56.e.01 offered poor protection in small animal models, future immunology efforts surrounding H5N1 vaccines may benefit from explicit avoidance of antigenic exposure to the head-stem interface epitope, which may be exposed by denaturation or re-organization of the protein during inactivation.^69^ Cleavage of HA, occurring during or after viral budding,^70^ has been shown to stabilize its closed trimeric conformation,^71^ decreasing accessibility of other protomer interface epitopes.^17,35^ Therefoore, stabilization and presentation of the mature, trimeric form of HA as it appears on virions may be a powerful strategy to minimize non-neutralizing epitope exposure to the immune system. One recent study found that H5 trimer-stabilizing mutations could improve the quality of vaccine-elicited antibody responses across multiple H5 immunogens.^72^ These closed-trimer immunogens elicited significantly higher neutralizing responses against both H5N1 and H1N1 strains, in part by shifting antigen recognition to the RBS, thereby reducing stimulation of responses to cryptic or nonessential epitopes.^72^ Similar studies involving stabilized versions of both H1 and H3 HA trimers, in which buried interface epitopes are shielded from exposure, have also reported superior immunogen performance.^73,74^ Ultimately, our understanding of influenza virus antigenicity is still incomplete. As new immunogens are tested and novel epitopes are unveiled, rigorous functional characterization is required to ensure a productive feedback loop.

## MATERIALS AND METHODS

### Cell line handling

Human embryonic kidney (HEK) cells derived from the Expi293F expression system (Thermo Fisher Scientific, Cat. No. A14527) were cultured at 37 °C with 8% CO_2_ in Expi293F expression media (Thermo Fisher Scientific). Cells were routinely passaged at mid-exponential phase (3.0×10^6^ cells/mL) in baffled shake flasks. Sf9 cells (*Spodoptera frugiperda* ovarian cells, ATCC, Cat. No. CRL-1711) were grown in Sf-900 II SFM medium (Thermo Fisher Scientific) supplemented with 100 U/mL of penicillin and 100 μg/mL of streptomycin (Thermo Fisher Scientific) at 27 °C with shaking in baffled flasks, or stationary at 27 °C in adherent T-175 flasks during baculoviral propagation. MDCK-SIAT1 cells (Madin-Darby canine kidney cells engineered for stable human α2,6-sialyltransferase expression, Sigma-Aldrich, Cat. No. 05071502) were grown in T-175 flasks at 37 °C with 5% CO_2_ in Dulbecco’s modified Eagle’s medium (DMEM) enriched with high glucose (Thermo Fisher Scientific), along with 10% heat-inactivated fetal bovine serum (FBS, Thermo Fisher Scientific), 1× 4-(2-hydroxyethyl)-1-piperazineethanesulfonic acid (HEPES, Gibco), and 1× GlutaMax (Thermo Fisher Scientific). MDCK-SIAT1 cells were routinely passaged at confluency using a 0.25% trypsin-EDTA solution (Thermo Fisher Scientific) for cell detachment.

### Influenza A virus

H1N1 A/California/04/2009 viruses were propagated in MDCK-SIAT1 cells and harvested at 72 h post-infection. TCID_50_ (tissue culture infectious dose 50) was used to quantify the viral titer. Briefly, serial dilutions of the virus sample were made and added to MDCK-SIAT1 cells supplemented with 1 μg/mL TPCK-trypsin. After incubation for 72 h, the cells were examined for cytopathic effect (CPE). The dilution that caused infection in 50% of the wells was used to calculate the TCID_50_. The Reed-Muench method was used to estimate the viral titer in terms of TCID_50_/mL. Mouse-adapted H3N2 537 A/Philippines/2/1982 (X-79, 6:2 A/PR/8/34 reassortant) virus was grown in 10-day-old embryonated chicken eggs at 37 °C for 48 h and was cooled at 4 °C overnight. Cell debris was removed by centrifugation at 4000 ×g for 20 min at 4 °C. Virus aliquots were stored at −80 °C until use.

The influenza A/Texas/37/2024 human H5N1 eight-plasmid reverse genetics system was used to construct the virus.^75^ In brief, eight DNA plasmids were cloned into a pHW2000 vector and transfected into cocultured MDCK-SIAT1 cells and HEK 293T cells. After 72 h, the supernatants were collected. Viruses were plaque purified on MDCK-SIAT1 cells grown in DMEM (Gibco) containing 10% FBS (Gibco) and penicillin–streptomycin mix (Gibco). Individual plaques were picked and grown in fresh MDCK-SIAT1 cells. To confirm the HA and NA sequences of the virus, viral RNAs were extracted from the supernatant and HA and NA segments were amplified and confirmed by Sanger sequencing.

### Recombinant IgG/Fab expression and purification

Synthetic DNA encoding VH/VL genes of antibodies (IDT) were cloned into a pCMV3 plasmid backbone (Fabs, human IgGs) or a pVRC plasmid backbone (mouse IgGs) using Gibson assembly and transformed into *E*.*coli* DH5α. Clones containing the correct plasmid sequences were used to inoculate 50 mL LB cultures, which were grown at 37 °C for 18 h before the plasmid DNA was harvested by midiprep (ZymoPURE II Plasmid Midiprep Kit). Plasmids (containing an N-terminal mouse kappa signal peptide secretion sequence) were then used to transfect Expi293F cells (3.0×10^6^ cells/mL density) at a 1:2 molar ratio of light:heavy chain IgG/Fab DNA using Expifectamine transfection kit (Thermo Fisher Scientific) according to manufacturer’s directions. 18 h after transfection, Transfection Enhancers 1 and 2 (Thermo Fisher Scientific) were added to expression cultures to increase yield. Expressions were allowed to proceed for 6 days at 37 °C / 8% CO_2_, after which cells were pelleted by centrifugation at 4,500 rcf for 50 min. The resulting supernatants were sterilized via 0.22 μm vacuum filtration (Sigma) and added to CaptureSelect CH1-XL beads (Fabs, human IgGs) (Thermo Fisher Scientific; 1 mL beads / 200 mL expression volume) or CaptureSelect IgG-Fc Multispecies affinity resin (mouse IgGs) (Thermo Fisher Scientific, 2 mL beads / 200 mL expression volume). The beads and supernatants were rotated for at least 2 h at 4 °C, then washed with 1× phosphate buffered saline (PBS) prior to three sequential elutions with 60 mM sodium acetate, pH 4.0 (CH1-XL) or 100 mM Glycine, pH 3.0 (IgG-Fc Multispecies). Eluants were pooled and neutralized with 1 M Tris, pH 8.0, and then buffer-exchanged four times into PBS using 30 kDa molecular weight cutoff (MWCO) ultrafiltration concentrators (Millipore). Protein concentration was then determined using a Nanodrop One (Thermo Fisher Scientific) using the respective theoretical extinction coefficient (calculated using ExPasy ProtParam) at 280 nm. IgGs and Fabs were stored at 4 °C until use.

### Recombinant Hemagglutinin (HA) expression and purification

Synthetic genes encoding the HA ectodomain amino acid sequence (IDT, Addgene) were subcloned into a customized pFastBac baculoviral expression vector using Gibson assembly. This vector contained an N-terminal gp67 signal peptide secretion sequence, along with a C-terminal thrombin protease cleavage site followed by a foldon trimerization domain, an AviTag/biotin ligase BirA recognition sequence, and a 6xHis tag. For the expression of the truncated HA head constructs, the trimerization domain and AviTag were omitted. Clones were transformed into *E. coli* DH5α, and plasmid DNA from colonies displaying the correct sequence were harvested by miniprep (NEB plasmid miniprep kit) and transformed into *E. coli* DH10Bac (Thermo Fisher Scientific, containing the Bac-to-Bac baculoviral expression system). Colonies displaying successful recombination were picked by blue/white screening and miniprepped to harvest purified baculoviral DNA (PureLink HiPure Plasmid Miniprep kit, Thermo Fisher Scientific). Sf9 cells (1.0×10^6^/mL, 10 mL) were transfected with baculoviral DNA using Cellfectin transfection reagent (Thermo Fisher Scientific) according to manufacturer’s protocols. Sf9 culture supernatant was harvested after 5 d of incubation at 27 °C and used to infect 50 mL of fresh Sf9 cells at a multiplicity of infection (MOI) of 1.0 for baculovirus amplification. After 5 d, baculovirus-containing supernatant was again harvested, clarified, and sterilized. This passaged baculovirus was then used to infect large-scale expression cultures of Sf9 cells (1.0 × 10^6^/mL, 1 L) at an MOI of 1.0. Expression cultures were incubated in baffled flasks at 27 °C with shaking for 3 d, then the cells were pelleted at 4,000 rcf for 20 min and the supernatant was collected. Ni Sepharose Excel resin (Cytiva, 2.5 mL settled resin / L culture) was added to protein-containing supernatants and binding proceeded overnight at 4 °C with rotation. The following day, the bound resin was collected via gravity-flow columns, washed sequentially with wash buffer 1 (20 mM sodium phosphate, 500 mM NaCl, 20 mM imidazole, pH 7.4) and wash buffer 2 (20 mM sodium phosphate, 500 mM NaCl, 40 mM imidazole, pH 7.4) prior to elution with elution buffer (20 mM sodium phosphate, 500 mM NaCl, 250 mM imidazole, pH 7.4). Elution was performed thrice, and all eluants were pooled and concentrated to 1.0 mL in 30 kDa molecular weight cutoff (MWCO) ultrafiltration concentrators (Millipore), filtered to remove precipitates, then further purified via size-exclusion chromatography (SEC). SEC was performed on a HiLoad 16/100 Superdex 200 prep grade column (Cytiva) in 20 mM Tris-HCl (pH 8.0) with 100 mM NaCl. Fractions containing properly folded HA proteins were collected, concentrated again, and quantified using a Nanodrop One (Thermo Fisher Scientific), using the respective theoretical extinction coefficient (calculated using ExPasy ProtParam) at 280 nm. HA proteins were aliquoted, flash-frozen, and stored at −80 °C until use.

### Antibody binding determination using ELISA

The enzyme-linked immunosorbent assay (ELISA) was performed to determine monoclonal antibody binding efficiency. HA proteins were diluted to 1 μg/mL in 1× PBS and used to coat Nunc MaxiSorp plates (Thermo Fisher Scientific) overnight at 4 °C (100 ng HA/well). The following day, the plates were dumped and washed thrice in PBS supplemented with 0.1% v/v Tween-20 (PBS-T), then blocked using a solution of 5% nonfat dry milk powder in ddH_2_O for 2 h at room temperature. The plates were again washed in PBS-T, then purified IgGs were diluted in PBS and added to the plates at a 10 μg/mL concentration. After 2 h equilibration at 37 °C, plates were emptied and washed thrice with PBS-T before adding horseradish peroxidase (HRP)-conjugated goat anti-human IgG antibody (Thermo Fisher Scientific) at a 1:5,000 dilution in PBS for 1 h at 37 °C. Plates were emptied and again washed 6 more times in PBS-T prior to adding 100 μL of 1-Step TMB Ultra ELISA Substrate Solution (Thermo Fisher Scientific) to each well. After a 10 min incubation in the dark at room temperature, reactions were quenched using 50 μL 2 M H_2_SO_4,_ and absorbance values were immediately measured at 450 nm (OD_450_) using a BioTek Synergy HTX Multimode Reader (Agilent). ELISAs were performed in duplicate for each antibody / HA pair, along with no-antibody and no-antigen negative controls to ensure assay rigor.

### Biolayer interferometry (BLI) assessment of binding interactions

BLI was conducted using an Octet Red96e instrument (Sartorius). For all experiments, 1× kinetics buffer (1× PBS at pH 7.4, and 0.002% v/v Tween 20) was used. For the measurement of K_d_, purified His-tagged HA protein (20 μg/mL) in 1× kinetics buffer was loaded onto HIS1K biosensors and incubated with purified Fabs at concentrations of 300 nM, 100 nM, and 33.3 nM. The assay protocol consisted of five steps: baseline (60 s in 1× kinetics buffer), loading (300 s with His-tagged HA protein), a second baseline (60 s in 1× kinetics buffer), association (120 s with Fabs), and dissociation (120 s in 1× kinetics buffer). K_d_ values were estimated using a 1:1 binding model. To determine binding competition, a His-tagged H1 A/California/2009 (H1N1) monomeric head construct was used to ensure all epitopes were equally accessible and prevent steric occlusion from HA trimer association. HA head (500 nM in 1× kinetics buffer) was first loaded onto HIS1K biosensors for 300 s. After a 60 s baseline, the probes were then saturated with competing Fab by exposing the sensors to 200 nM of relevant Fab in 1× kinetics buffer for 180 s, or until saturation was reached. The degree of additional binding was then assessed by immediately exposing the sensors to 200 nM of a second Fab in the presence of the first Fab (200 nM) for another 120 s. Each assay was run with a no-preloading/competing Fab control to determine the binding response of the second Fab in the absence of competition for comparison.

The competition index (CI) for the second Fab (Ab2) competing with a pre-bound Fab (Ab1) was calculated by:

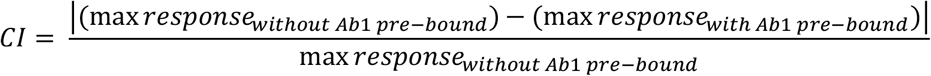

where “*max response*_*without Ab1 pre-bound*_” is the maximum response of Ab2 in the absence of Ab1 pre-bound and “*max response*_with Ab1 pre-bound_” is the maximum response of Ab2 in the presence of Ab1 pre-bound.

To measure binding to different forms of HA, Fabs were loaded onto a FAB2G probe until saturation, held for a 60 s baseline in kinetics buffer, then dipped into 300 nM of recombinant HAs for a 120 s association, then moved to kinetics buffer for a 120 s dissociation phase. Purified HA0 was cleaved via incubation with trypsin protease (TPCK-treated, Sigma) for 30 min at 37 ° C at a 50:1 substrate: enzyme ratio in HA buffer supplemented with 10 mM beta-mercaptoethanol (β-ME) to reduce the HA1/HA2 interchain disulfide. For the acid disassociation of HA, the purified protein was diluted 5-fold in 50 mM NaOAc buffer, pH 5.0 and incubated for 15 min at room temperature before dilution in kinetics buffer for binding measurements.

### Cryo-EM sample preparation, data collection, and data processing

The sequence of A/American black duck/New Brunswick/00464/2010 (H4N6) HA head from the S8V1-157 structure (PDB 8US0)^41^ was aligned with the HA ectodomain sequence of A/Darwin/9/2021 (H3N2) to create the truncated H3/Darwin21 head construct used for structure determination. The purified H3/Darwin21 head construct (250 μg) was combined in a 1:1:1:1 molar ratio of 01.z.01 Fab, ADI-85647 Fab, and FluA-20 Fab in 20 mM Tris-HCl (pH 8.0) with 100 mM NaCl. After complex formation for 1 h at room temperature, the proteins were filter-sterilized and injected onto a HiLoad 16/100 Superdex 200 preparative SEC column (Cytiva) in 20 mM Tris-HCl (pH 8.0) with 100 mM NaCl. The peak fractions corresponding to the relevant 3 Fab + 1 HA mass were collected and concentrated in spin filters (Millipore) to 1.8 mg/mL for cryo-EM sample preparation. In the case of 56.e.01 Fab, H3 head was similarly complexed with FluA-20 Fab and ADI-85647 Fab prior to SEC purification, after which 56.e.01 Fab was added in a 1:1 molar ratio to the HA + 2 Fab SEC peak and incubated for 1 h at room temperature. Protein complexation was manually inspected by running all SEC fractions on 12% acrylamide SDS-PAGE to confirm Fab-HA coelution. Due to the tendency of the 3Fab-H3 complexes to precipitate out of solution, EM grids were prepared immediately after SEC purification. n-octyl-β-d-glucoside [0.1% w/v] was added to the sample seconds before application (3.0 μL) to a 300-mesh Quantifoil R1.2/1.3 Cu grid pretreated via glow discharge. Excess liquid was blotted away using filter paper (SubAngstrom) with a blotting force of −1 and a blotting time of 2 s. The grid was then plunge-frozen in liquid ethane using an FEI Vitrobot Mark IV (Thermo Fisher Scientific). For the 01.z.01 complex, the clipped grid was loaded onto a Titan Krios microscope equipped with a Gatan BioQuantum K3 imaging filter and camera (Thermo Fisher Scientific). 4,000 movies were collected at an 81,000X nominal magnification, corresponding to a 0.529 Å pixel size. An accelerating voltage of 300 kV was applied in a total dose of 57.35 e^-^/Å^2^ split across 40 frames. A defocus range of −0.6 μm to −2.0 μm and a spherical aberration of 2.7 mm were applied to data collection. Movies were collected near the edges of holes to account for particle ice preference. For the 56.e.01 complex, the clipped grid was loaded onto a Glacios 2 Cryo-TEM, equipped with a Falcon 4i Direct Electron Detector. 280 movies were recorded at a 150,000× nominal magnification, corresponding to a 0.96 Å pixel size. An accelerating voltage of 200 kV was applied in a total dose of 60 e-/Å^2^. A defocus range of −0.3 μm to −3 μm and a spherical aberration of 2.7 mm were applied to data collection.

CryoSPARC^76^ was used to process raw movies, pick particles, and refine an initial 3D volume map of the complex, which was sharpened using DeepEMhancer v0.15.^77^ For a more detailed processing workflow, refer to **Figure S1**. The 01.z.01 Fab + H3/Darwin21 complex was generated using AlphaFold 3^56^ and fitted into the density of the complex map using UCSF ChimeraX.^78^ Coot^79^ was then used to manually adjust the model in an iterative refinement cycle alongside Phenix real-space refinement program.^80^ This process was repeated until no progress was observed, and the final model was then validated using the PDB online validation server equipped with MolProbity. Further structural analysis of the model and figure generation were performed in UCSF ChimeraX.

### Hemagglutination activity inhibition (HAI) assay

4 HA units (HAU) in 50 µL of either H3N2 A/Philippines/2/1982 or H1N1 A/California/04/2009 virus was added to 2-fold serially diluted antibodies (starting at 100 μg/mL) in 50 µL PBS in a V-bottom 96-well plate. After a 1 h equilibration at room temperature, 50 μL of gently resuspended 1% turkey red blood cells (Innova) was added to the wells. After 20 min, hemagglutination was observed, and the highest dilution of the antibody that prevented hemagglutination was recorded as the HAI titer. HAI assays were repeated in duplicate.

### Microneutralization assay

MDCK-SIAT1 cells were seeded in 96-well plates and grown to 100% confluency. Cells were washed in warm PBS to remove media, then minimal essential media (MEM) (Gibco) containing 25 mM HEPES (Gibco) was added to the cells. Purified antibodies were 2-fold serially diluted (starting at 100 μg/mL) in MEM with 25 mM HEPES, then 100 TCID_50_ (median tissue culture infectious dose) of respective viruses were added, and the mixture was incubated at 37 °C for 1 h. The virus-antibody mixture was then added to the cells, and infection proceeded for 1 h at 37 °C. Afterwards, virus-containing media was discarded, and fresh MEM supplemented with 25 mM HEPES and TPCK-trypsin (Sigma Aldrich, 1 μg/mL final concentration) was added. This media also contained the same serial dilution of antibodies as before to ensure assay detection of viral neutralization mechanisms beyond initial receptor binding inhibition. After 72 h at 37 °C, virus presence was detected using the HAI assay as described above to determine antibody median neutralizing concentration (MN_50_).

### Prophylactic protection experiments and lung titer determination

Female BALB/c mice at 6 weeks old (n = 5 per group) were anesthetized with isoflurane and dosed with 200 μg (10 mg/kg) of purified mouse-adapted antibodies (mIgG: 1 or 2c subtype, injected intraperitoneally). 4 h after antibody administration, the mice were again anesthetized with isoflurane and intranasally administered a 5x median lethal dose (LD_50_) of either mouse-adapted H3N2 A/Philippines/2/1982 (X-79, 6:2 A/PR/8/34 reassortant) or human clinical isolate H1N1 A/California/04/2009 virus (H1/Cal09). Weight loss was monitored daily over the 14-day course of the experiment, with the humane endpoint defined as a weight loss of 25% from the starting weight on day 0. A positive control group was included using 5 mg/kg CR9114^15^ mAb, while a negative control group was administered only PBS during antibody injection.

To determine lung titer, mice in separate groups (n = 3 per group) received the above treatment, except they were sacrificed 72 h post-infection. Lungs of infected mice were surgically extracted and homogenized in 1 mL of MEM with 1 μg/mL of TPCK-trypsin using a gentleMACS Dissociator (Miltenyi Biotec). Subsequently, viral lung titers were measured by TCID_50_ (median tissue culture infectious dose) assay. Briefly, serial log_2_ dilutions of lung homogenates were added to confluent MDCK-SIAT1 cells seeded in 96-well plates and incubated for 72 h at 37 °C. Cells were individually inspected under a light microscope for cytopathic effect and the dilution that caused infection in 50% of the wells was used to calculate the TCID_50_. The Reed-Muench method was used to average the viral titer across 3 technical assay replicates per mouse.

All H1N1 and H3N2 animal experiments were performed in a BSL-2 facility in accordance with the protocols approved by UIUC’s Institutional Animal Care and Use Committee (IACUC).

*In vivo* H5N1 protection experiments were conducted in The Chinese University of Hong Kong Biosafety Level 3 (BSL-3) facility. Per experimental group, a total of 8 female BALB/c mice at 6 weeks old were anesthetized with isoflurane and intranasally infected with 5× LD_50_ of wild-type A/Texas/37/2024 (H5N1) virus. Mice were given the indicated antibody at a dose of 25 mg/kg intraperitoneally at 4 h before infection. Weight loss was monitored daily for 14 days in 5 out of 8 mice per experimental group. The humane endpoint was defined as a weight loss of 25% from initial weight on day 0. To determine the lung viral titers on day 3 post-infection, lungs were harvested from the remaining three mice in each experimental group and homogenized in 1 mL of MEM with 1 μg/mL of TPCK-trypsin using a gentleMACS Dissociator (Miltenyi Biotec). Subsequently, viral titers were measured by TCID_50_ assay. The study protocol was carried out in strict accordance with the recommendations and was approved by the Animal Experimentation Ethics Committee of the Chinese University of Hong Kong (25-190-HMF).

### Multiple sequence alignments

All amino acid sequences for influenza HAs were obtained from GenBank or the GISAID EpiFlu database. Alignments were performed by MAFFT using G-INS-i algorithm.^81^ The resulting clustal files were visualized using ESPript3.2.^82^ Antibody germline gene assignment and alignments were calculated using IMGT/V-Quest.^83^

## DATA AVAILABILITY

Cryo-EM maps have been deposited in the Electron Microscopy Data Bank (EMDB) with accession numbers: EMD-79286 (01.z.01-H3 head) and EMD-79287 (56.e.01-H3 head). The corresponding refined model of the 01.z.01-H3 head complex has been deposited in the Protein Data Bank (PDB) with accession number 39AW.

## ACKNOWLEDGEMENTS

We thank Dr. Kristen Rhebergen, Dr. Tiit Lukk, and the Materials Research Laboratory Central Research Facilities at the University of Illinois Urbana-Champaign, as well as Dr. Frank Vago and the Cryo-EM Facility at Purdue University, for access to cryo-EM instrumentation. We thank Professor Kevin McCarthy for his helpful comments and for generously supplying expression constructs for S8V1-157. We also thank Professor Florian Krammer for providing the H3N2 A/Philippines/2/1982 (X-79) virus and Dr. Qi Wen Teo for providing the H1N1 A/California/04/2009 virus. Finally, we thank Dr. Wenhao O. Ouyang for insightful discussions.

## AUTHOR CONTRIBUTIONS

G.L.S. and N.C.W. conceived of and designed the study. G.L.S., H.L., T.P. and C.Z. performed the experiments. C.C., Y.S., and S.T.T. performed the H5N1 experiments under the supervision of C.K.P.M. G.L.S. was supervised by D.A.M. and N.C.W. G.L.S. drafted the paper with editorial and revisional oversight from N.C.W. All authors read and revised the manuscript.

