## Supplementary material for "Antibodies targeting a conserved cryptic epitope at the influenza hemagglutinin head-stem interface via distinct binding modes": Supplmentary Information

**Table S1. Cryo-EM data collection, refinement and validation statistics.**

| | 01.z.01 Fab + Dar21 HA head + FluA-20 Fab + ADI-85647 Fab (EMBD-XXXXX) (PDB XXXX) | 56.e.01 Fab + Dar21 HA head + FluA-20 Fab + ADI-85647 Fab (EMBD-XXXXX) |
|---|---|---|
| **Data collection and processing** | | |
| Magnification | 81,000 | 150,000 |
| Voltage (kV) | 300 | 200 |
| Electron exposure ($e^-$/Å$^2$) | 57.35 | 60.00 |
| Defocus range (μm) | -0.6 – 2.0 | -0.3 – 3.0 |
| Pixel size (Å) | 0.529 | 0.96 |
| Symmetry imposed | C1 | C1 |
| Initial particle images (no.) | 407,616 | 71,910 |
| Final particle images (no.) | 103,663 | 4,903 |
| Map resolution (Å) | 3.38 | 8.77 |
| FSC threshold 0.143 | | |
| Map postprocessing | DeepEMhancer | N/A |
| **Refinement** | | |
| Initial model used (PDB code) | N/A | N/A |
| Model composition | | |
| Non-hydrogen atoms | 3,998 | |
| Protein residues | 517 | |
| Root-mean-square deviations | | |
| Bond lengths (Å) | 0.005 | |
| Bond angles (°) | 0.824 | |
| Validation | | |
| MolProbity score | 1.99 | |
| Clashscore | 9.63 | |
| Poor rotamers (%) | 2.01 | |
| Ramachandran plot | | |
| Favored (%) | 96.28 | |
| Allowed (%) | 3.52 | |
| Disallowed (%) | 0.20 | |

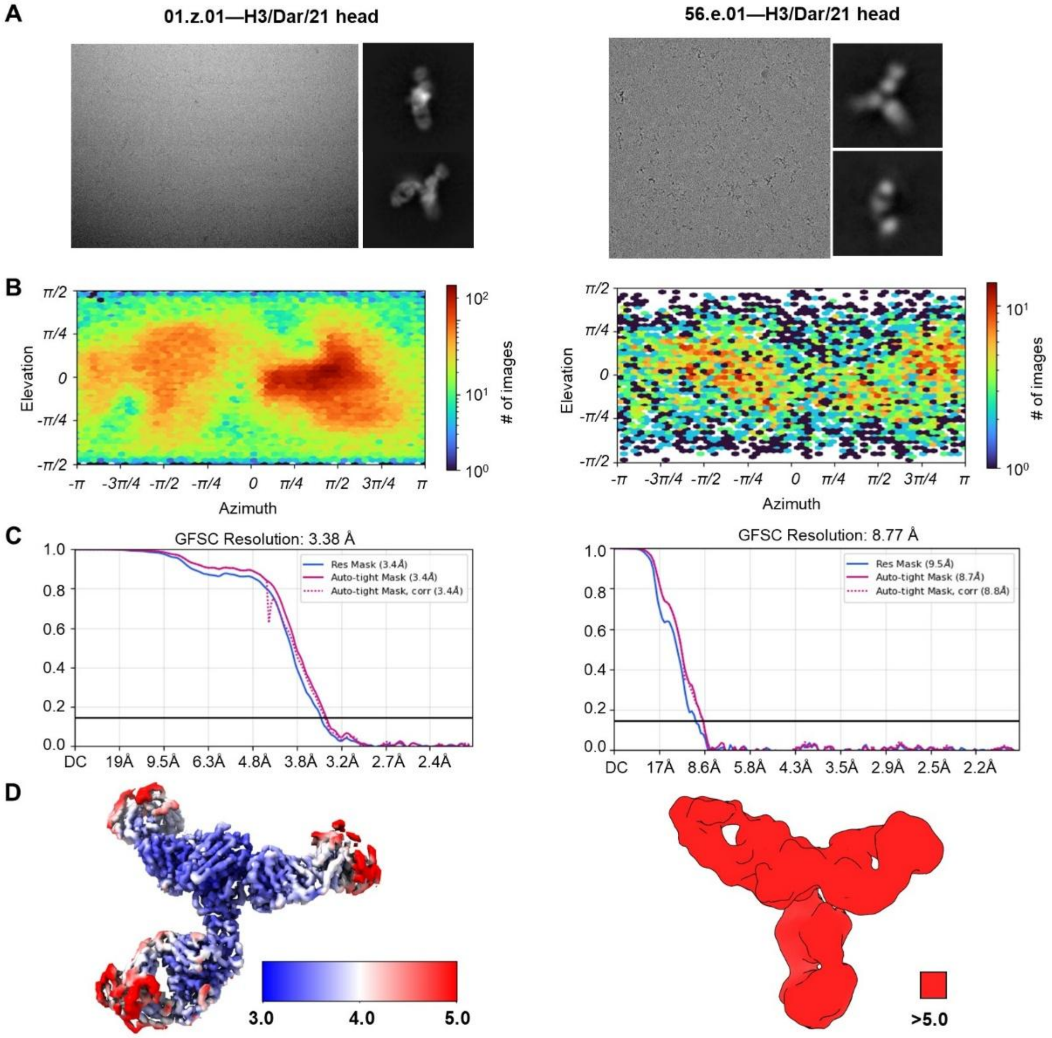


**Figure S1: Cryo-EM data processing of Fab-HA complexes**

CryoSPARC[1] data processing is shown for the 01.z.01 Fab + A/Darwin/9/2021(H3N2) HA head construct (left) and the 56.e.01 Fab + A/Darwin/9/2021(H3N2) HA head construct (right).

**(A)** Representative micrographs and 2D class averages for selected particles. 01.z.01 + H3 HA: 3820 micrographs, 407,616 particles picked, 103,663 particles selected to build 3D map. 56.e.01 + H3 HA: 290 micrographs, 71,910 particles picked, 4,903 particles selected to build 3D map.

**(B)** Viewing direction distribution of selected particles used to build each map.

**(C)** Gold-standard Fourier shell correlation curves for each map, with the 0.143 cutoff marked by a horizontal black line.

**(D)** Local resolution analysis of the respective complex maps.

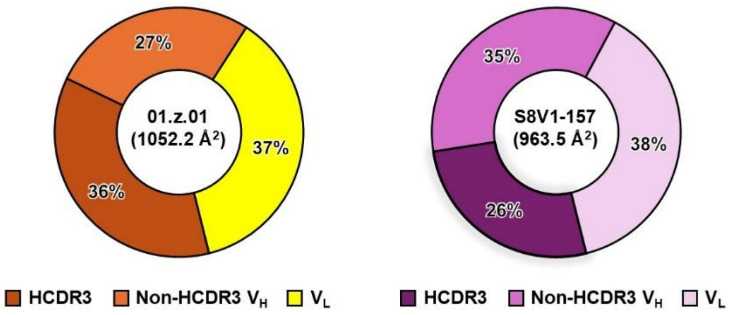


**Figure S2. Buried surface area (BSA) analysis for head-stem interface antibodies**

The paratope BSA for 01.z.01 is reported (left) as a fraction of the HCDR3 (dark orange), non-HCDR3 $V_H$ (light orange), and $V_L$ (yellow). The paratope BSA is similarly reported for S8V1-157 (PDB 8US0,[2] right) as a fraction of the HCDR3 (purple), non-HCDR3 $V_H$ (orchid), and $V_L$ (pink). Paratope BSA was calculated by PDBePISA[3].

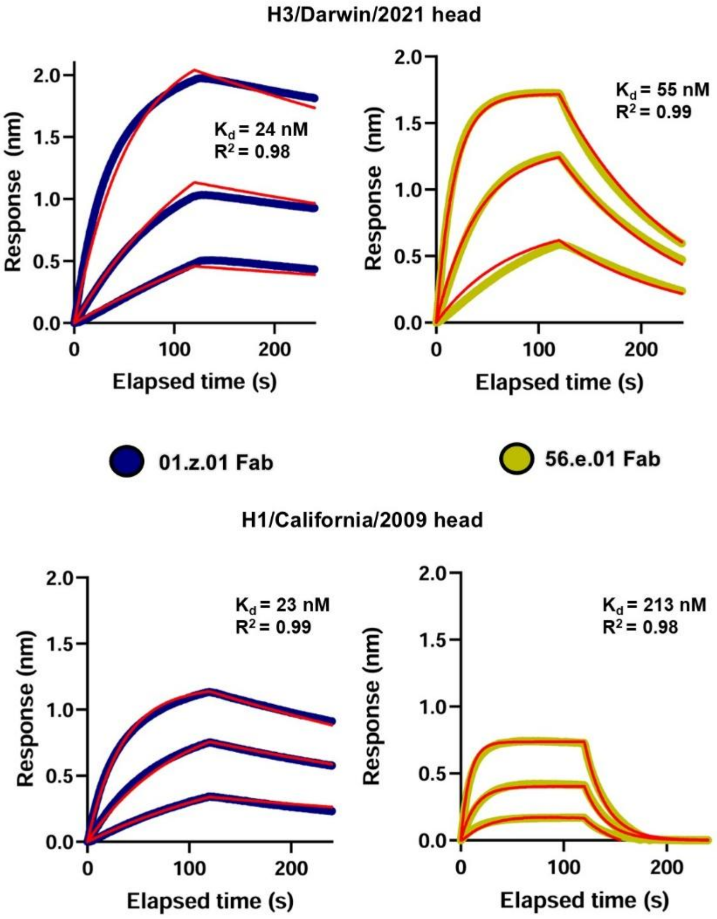


**Figure S3. Kinetic profiling of head-stem interface Fab binding to HA head domain**

Biolayer interferometry (BLI) data were collected at three Fab concentrations (300 nM, 100 nM, and 33 nM) on either immobilized A/Darwin/9/2021 (H3N2) HA1 (top) or A/California/04/2009 (H1N1) HA1 (bottom). Binding curves are shown for 01.z.01 Fab (left, dark blue) and 56.e.01 Fab (right, yellow). Curve fitting using a 1:1 binding model (red lines) was used to calculate dissociation constants ($K_d$).

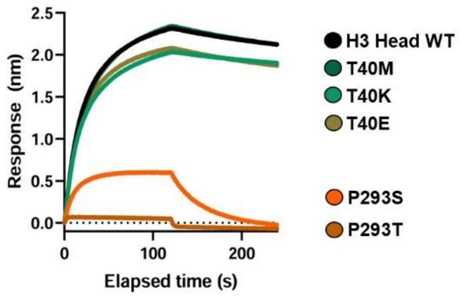


**Figure S4. Binding analysis of S8V1-157 on H3 head mutants**

Biolayer interferometry binding curves for 500 nM S8V1-157 Fab[2] against a panel of immobilized A/Darwin/9/2021 (H3N2) HA1 single-site variants.

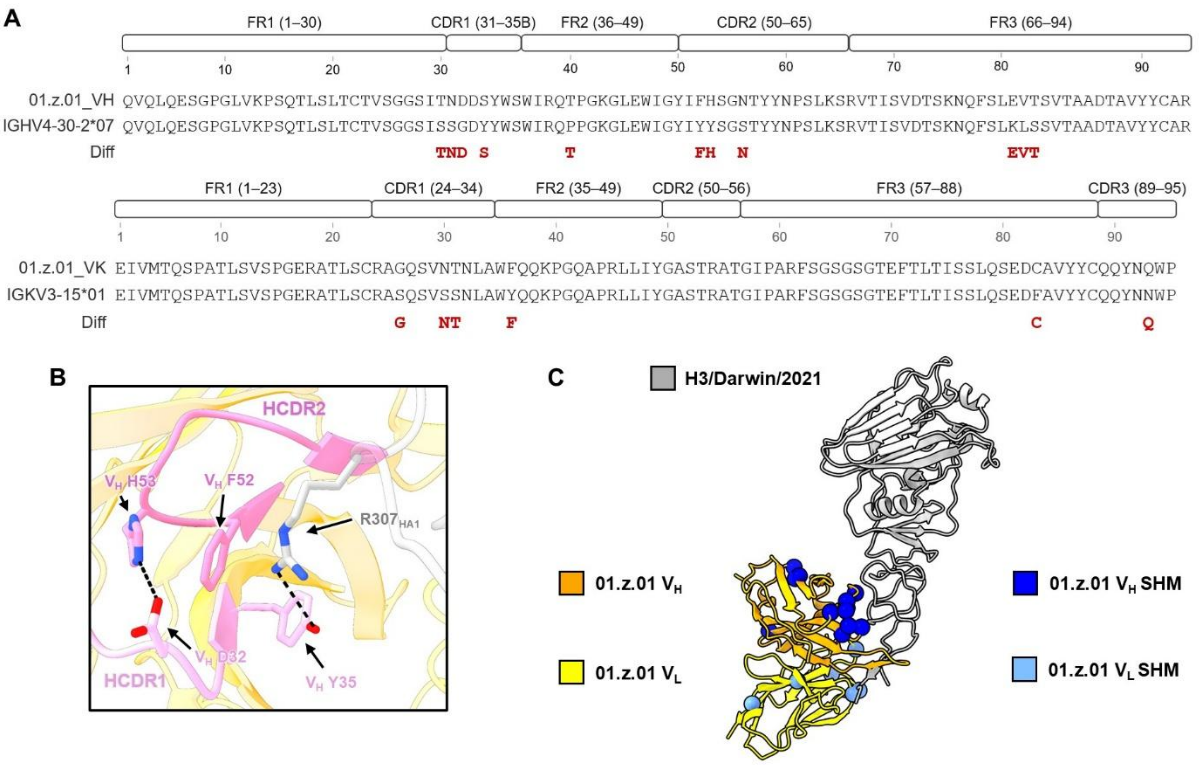


**Figure S5. Germline gene alignment and somatic hypermutation analysis of 01.z.01**

**(A)** Sequence alignment of 01.z.01 $V_H$ (top) and $V_L$ (bottom) with their respective inferred germline genes. Kabat numbering is employed. Locations of somatic hypermutations (SHMs) are shown in red.

**(B)** Molecular depiction of the involvement of residues arising from 01.z.01 SHM with the highly conserved $R307_{HA1}$. For clarity, the 01.z.01 HCDR1 (light pink) and HCDR2 (pink) are labeled.

**(C)** Ribbon diagram of 01.z.01 scFv ($V_H$: orange, $V_L$: yellow) in complex with A/Darwin/9/2021 (H3N2) HA1 (light gray). $C_\alpha$ atoms from residues arising from 01.z.01 SHM are depicted as spheres ($V_H$ SHM: dark blue, $V_L$ SHM: light blue).
